# Exploring variation in human papillomavirus vaccination uptake: multi-level spatial analysis

**DOI:** 10.1101/210260

**Authors:** Maurane Riesen, Garyfallos Konstantinoudis, Phung Lang, Nicola Low, Christoph Hatz, Mirjam Maeusezahl, Anne Spaar, Marc Bühlmann, Ben D. Spycher, Christian L. Althaus

## Abstract

**Background:** Understanding the factors that influence human papillomavirus (HPV) vaccination uptake is critically important to design effective vaccination programmes. In Switzerland, completed HPV vaccination by age 16 years among women ranges from 30 to 79% across 26 cantons (states). Our objective was to identify factors that are associated with the spatial variation in HPV vaccination uptake.

**Methods and findings:** We used data from the Swiss National Vaccination Coverage Survey 2009-2016 on HPV vaccination status (≥1 dose) of 14-17 year old girls, their municipality of residence and their nationality for 21 of 26 cantons (N=8,965). We examined covariates at municipality level: language, degree of urbanisation, socio-economic position, religious denomination, results of a vote about vaccination laws; and, at cantonal level, availability of school-based vaccination and survey period. We used a series of conditional auto regressive (CAR) models to assess the effects of covariates while accounting for variability between cantons and municipal-level spatial autocorrelation. In the best-fit model, school-based vaccination (adjusted odds ratio, OR: 2.51, 95% credible interval, CI: 1.77-3.56) was associated with increased uptake, while lower acceptance of vaccination laws was associated with lower HPV vaccination uptake (OR 0.61, 95% CI: 0.50-0.73). Overall, the covariates explained 88% of the municipal-level variation in uptake.

**Conclusions:** In Switzerland, cantons play a prominent role in the variation in HPV vaccination uptake, especially through the provision of school-based vaccination delivery. HPV vaccination uptake is also strongly associated with inhabitants’ attitudes towards vaccination. To increase uptake, efforts should be made both to mitigate vaccination scepticism and to encourage school-based vaccination.

## Introduction

Human papillomavirus (HPV) is the most common viral infections of the reproductive tract [1]. Persistent infections with HPV types 16 and 18 are responsible for 70% of cervical cancers and precancerous cervical lesions [1]. Genital HPV types also cause anogenital warts and cancers of the anus, vulva, vagina and penis [1]. In 2006, the first vaccine against HPV was licensed and, by 2016, at least 68 countries had implemented vaccination programmes for the prevention of cervical cancer in at least one region [2]. Optimal HPV vaccination coverage is estimated to be around 70% for women but there are large geographical disparities in vaccination coverage between and within countries [2,3]. The United Kingdom (UK) and Australia have reached homogeneous levels of around 70% vaccination coverage [4–6]. In contrast, many countries including Italy, France, Switzerland, Germany, the Netherlands and the United States (US) experience lower national coverage rates with large regional variations [7–11].

HPV vaccine is seen as a challenge for achieving high levels of coverage for several reasons, including concerns that it might promote risky sexual behaviour in adolescents and logistical issues about reaching adolescents [12]. Individuals shape the geographic and social contexts in which they live, but their behaviour is also affected by their context [13]. An understanding of community and individual level factors associated with HPV vaccination uptake is therefore important. It has been shown that countries with extensive school-based vaccination reached markedly higher uptake rates [14]. Conversely, lower levels of HPV vaccine uptake have been found in communities or states with high levels of votes for religious or conservative parties [11][15], but it is not known how closely these reflect attitudes towards vaccination. At the individual level, findings about factors such as socio-economic position (SEP), ethnicity or religious affiliation are more mixed [8,11,16–22]. For example, poverty, based on either low income or SEP has been found to be associated with both lower [11,16,19,20,23–26] and higher HPV vaccination uptake [18,20,21,27]. Furthermore, few studies have accounted for spatial autocorrelation due to unmeasured confounding [16,17,22]. Neglecting this spatial autocorrelation can lead to spurious associations [28].

Switzerland provides a valuable setting for investigating regional differences in vaccine uptake. The country is spatially divided into 26 cantons (states) and four language regions (German, French, Italian and Romansh). The cantons have a high degree of autonomy with devolved administration of health and education. Within cantons, municipalities also enjoy a high level of autonomy including the power to levy taxes and pass municipal laws. People vote regularly in referendums on a wide range of issues that then determine legislation. All Swiss cantons implemented HPV vaccination programmes by the end of 2008, targeting 11 to 14 years old girls for basic vaccination and additionally including young women and men (up to 26 years old) for complementary vaccination. The HPV basic vaccination programmes for school-aged girls differ widely between cantons, ranging from the simple distribution of educational material, informing parents that vaccination is available, through to school-based vaccination delivery [29–31].

The objective of this study was to investigate the spatial heterogeneity of HPV vaccination uptake in Switzerland, and to identify factors at different spatial levels that explain this variation. We investigated both political and cultural contextual factors. We hypothesised that the canton of residence would represent an important contextual factor influencing whether or not an individual girl would receive HPV vaccination. We further expected that covariates at the level of the municipality, such as the degree of political scepticism about vaccination, socio-economic status, language, religion or the level of urbanisation could represent important contextual factors that play a role in explaining differences in uptake.

## Methods

We conducted a multi-level spatial analysis of the Swiss National Vaccination Coverage Survey (SNVCS) [32,33]. We used a series of Bayesian hierarchical logistic regression models that include spatial autocorrelation, a random effect to account for variability between the cantons, and several covariates.

### Individual level data

We used data from the SNVCS, which is a national cross-sectional survey that monitors immunization coverage of children and adolescents [32,33]. The Swiss Federal Office of Public Health (FOPH) mandates the Epidemiology, Biostatistics and Prevention Institute (EPBI, University of Zurich, Switzerland) to collate data in three-year cycles from all cantons in surveys organised either by EBPI or by the individual cantons. Three different sampling methods were used: cluster sampling (municipalities), simple random sampling, or information collection by school nurses (online supplementary Table S1). The methodology is described in detail elsewhere [33]. For cluster sampling and simple random sampling, the parents of 16 years old girls received an invitation (by email or by phone) and were asked to send a copy of the daughter’s vaccination card. In three cantons, this information was collected by school nurses in 14-16 year old girls. We used anonymised, individual-level information about HPV vaccination status (having received at least one dose of HPV vaccine), nationality (Swiss or non-Swiss, coded as 0 and 1 respectively) and municipality of residence. Missing individual information about nationality was replaced with the proportion of non-Swiss people in the subject’s municipality of residence based on the national census in 2013 [34].

### Covariates at municipality and cantonal level

At the cantonal level, we considered survey period (2008-2010, 2011-2013, 2014-2016, excluding 2008 because not all cantons had implemented their vaccination programme at the time) and availability of HPV vaccination delivery in at least one school in the canton (yes, no) [35]. At municipality-level we considered language region (French, German, or Italian) [36], majority religious denomination (≥50% catholic, ≥50% protestant or neither) [37], socio-economic position (mean Swiss-SEP, a neighbourhood-based measure of with lower values indicating lower socio-economic position [38,39]), and level of urbanisation (rural, semi-urban, urban, based on standard Swiss classifications [36]).

We also considered the municipality-level results of a popular referendum in 2013 as a proxy measure of vaccine scepticism [40,41]. The referendum was proposed by opponents of a revised national law about the control of epidemics, which included new recommendations about vaccination. We calculated the percentage of votes in favour of revision of the law for each municipality. We considered municipalities with a low percentage of people in favour of the revised law as having higher proportions of people who are sceptical about vaccination than municipalities that strongly favoured the revised law. For the continuous variables, referendum results and the Swiss-SEP index, we generated quartiles and compared the lowest and the highest quartiles with the second and third (baseline) quartiles to capture possible effects of more extreme values. A more detailed description of the variables is given in the online supplementary material (Section 1, Fig S1-S6 and Table S2).

The SNVCS has received ethical committee approval. According to the Swiss Human Research Act (HRA, Art.2.2 al.c.), additional ethical committee review for this study was not required because anonymised health-related data were used.

### Statistical analysis

We developed Bayesian hierarchical logistic regression models to investigate the spatial heterogeneity of HPV vaccination uptake across municipalities (Table 1). First, we fitted a model that captures spatial variation at the municipality level (model 1). Spatial autocorrelation was modelled using the Besag– York–Mollié (BYM) conditional autoregressive (CAR) prior distribution [42,43]. For municipalities not sampled by the survey, BYM borrows information about the uptake from the neighbouring municipalities. Second, we added a random effect at the cantonal level (model 2) to test the hypothesis that cantons represent an important contextual factor for HPV vaccination uptake. Third, we additionally included all covariates (model 3, ‘full model’) and calculated the percentage of municipal variation explained by the cantonal random effect and the covariates. To do this we calculated the median posterior variance of the municipal random effect (sum of spatially correlated and uncorrelated component) in each of these models and the percentage reduction of this variance in models 2 and 3 compared to model 1.

**Table 1.** Characteristics of the included participants of the Swiss National Vaccination Coverage Survey (SNVCS) on HPV.

| Covariates |  | Total<br>N | (%) | Vaccination uptake<br>(%) | N |
| --- | --- | --- | --- | --- | --- |
| Nationality | Unknown | 1051 | 0.12 | 0.66 | 690 |
|  | Swiss | 6254 | 0.70 | 0.51 | 3199 |
|  | Non-Swiss | 1660 | 0.19 | 0.60 | 991 |
| Urbanisation levels<sup>1</sup> | rural | 1549 | 0.17 | 0.49 | 758 |
|  | urban | 1650 | 0.18 | 0.53 | 868 |
|  | semi-urban | 5766 | 0.64 | 0.56 | 3254 |
| SEP quartile<sup>1</sup> | lowest SEP | 4839 | 0.54 | 0.53 | 2582 |
|  | baseline SEP | 2247 | 0.25 | 0.59 | 1323 |
|  | highest SEP | 1879 | 0.21 | 0.52 | 975 |
| Political opinion<sup>1</sup> | lowest acceptance | 5198 | 0.58 | 0.59 | 3055 |
|  | baseline acceptance | 2264 | 0.25 | 0.38 | 856 |
|  | highest acceptance | 1503 | 0.17 | 0.64 | 969 |
| Religious<br>demonation<sup>1</sup> | No majority | 4099 | 0.46 | 0.57 | 2326 |
|  | ≥ 50% Protestant | 973 | 0.11 | 0.45 | 435 |
|  | ≥ 50% Catholic | 3893 | 0.43 | 0.54 | 2119 |
| Language region<sup>1</sup> | German speaking | 5941 | 0.66 | 0.49 | 2930 |
|  | French speaking | 2437 | 0.27 | 0.68 | 1658 |
|  | Italian speaking | 587 | 0.07 | 0.5 | 292 |
| School-based<br>vaccination<sup>2</sup> | no | 2936 | 0.33 | 0.37 | 1084 |
|  | yes | 6029 | 0.67 | 0.63 | 3796 |
| Survey period<sup>2</sup> | 2009-2010 | 2064 | 0.23 | 0.46 | 945 |
|  | 2011-2013 | 3793 | 0.42 | 0.54 | 2039 |
|  | 2014-2016 | 3108 | 0.35 | 0.61 | 1896 |

<sup>1</sup> municipality level covariate
<sup>2</sup> Cantonal level covariate

To examine the effect of the pre-specified covariates, we performed model selection using the deviance information criterion (DIC) [44]. In addition to models 1-3, we ran four alternative models that only included the municipal random effect and the covariates (model 4), the cantonal random effect (model 5), the cantonal random effect and the covariates (model 6), and the covariates only (model 7). We also examined the univariable association with each of the eight covariates (model 8). We present the univariable associations (model 8), a fully adjusted logit model excluding random effects (model 7) and the model from 1-7 with the smallest DIC. We present results as median odds ratios (OR) with 95% credible intervals (CI).

We conducted three sensitivity analyses. First, we examined whether the non-respondents differ from respondents with respect to the covariates. Second, we compared results of the main analysis with models that assumed that all non-respondents were vaccinated, or that all non-respondents were not vaccinated. Third, we examined whether the survey sampling method affected the results by including sampling method as a covariate in the model with the smallest DIC.

Inference was performed using the Integrated Nested Laplace Approximation (INLA) for latent Gaussian models [45]. Further details about the different models, their implementation and the sensitivity analyses are provided in the online supplementary material (Sections 2 and 3).

## Results

We analysed data from 21 of 26 cantons (91.1% of the Swiss population, online supplementary Table S1).

### Data characteristics

We analysed data from 8,965 out of 14,106 sampled girls from the participating cantons. We excluded 2,056 individuals sampled in 2008, 3,072 individuals who did not respond, and 13 individuals with missing information about municipality of residence. The average response rate was 75.1% and ranged from 39.9% to 92.1% between cantons (online supplementary Fig. S7). Amongst the included participants, data on nationality were missing in 11.7% (1,051/8,965), concentrated in four cantons (online supplementary material, Section 1). The average vaccination uptake in survey participants from the 21 cantons and over all survey periods was 53.2% (95% confidence interval, CI: 46.8%-59.7%) but varied greatly between cantons (Fig 1). Seventy percent of included girls in the survey were Swiss, 66% and 27% lived in German- and French-speaking municipalities, respectively. Sixty seven percent of included girls lived in a canton where school-based vaccination was available in one or more schools (Table 1).

**Fig 1.**
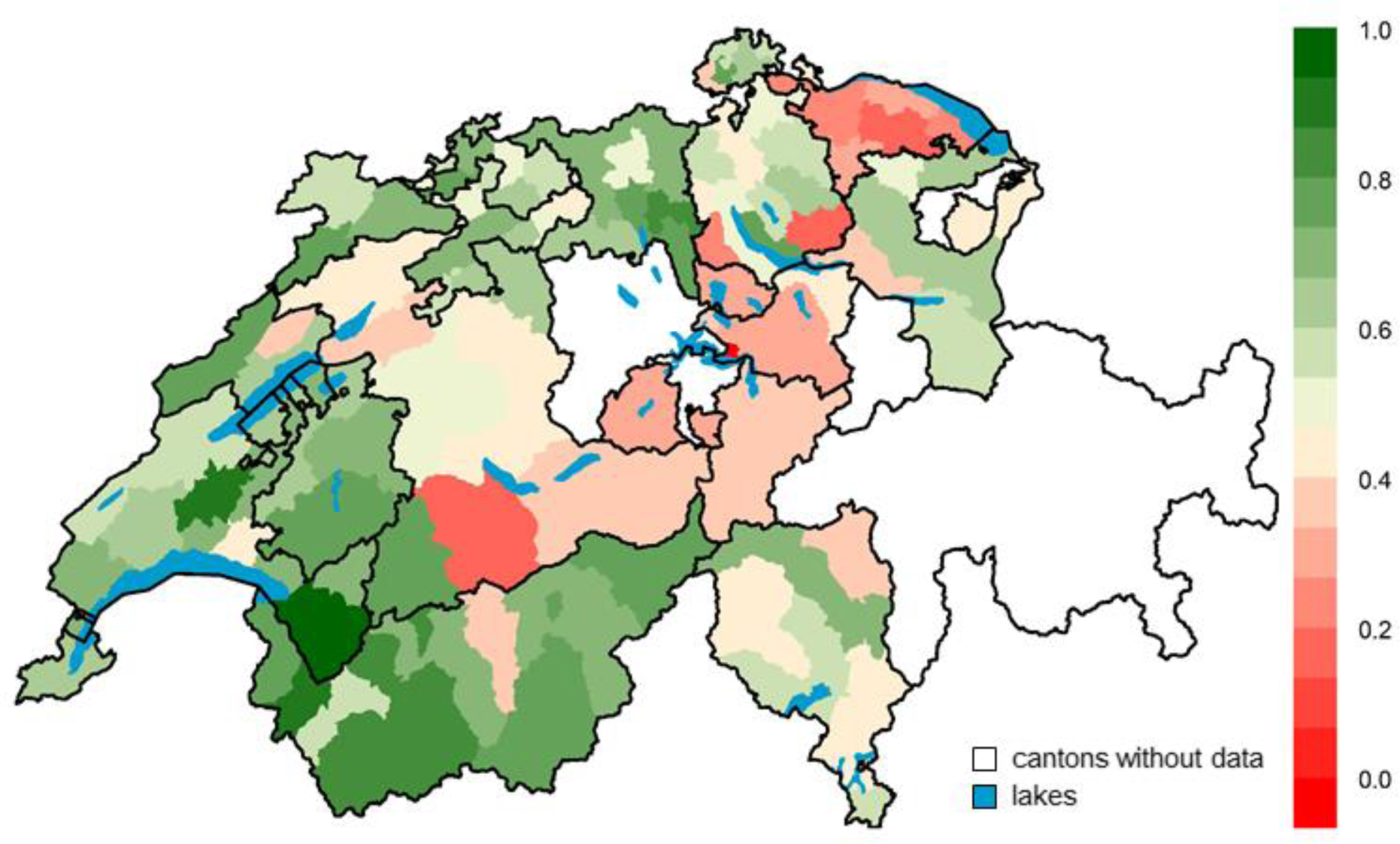
Crude HPV vaccination uptake per district in Switzerland over all survey periods (2009-2016). White areas represent cantons for which we did not get authorization to analyse the data.

### Spatial variability

Model 1 showed considerable spatial variation of HPV uptake at the municipal level (top panel, Fig 2). Including a random effect term at the cantonal level (Model 2) showed that about 63% of this variation is explained by cantonal differences (middle panel, Fig 2). Additionally including covariates further accounted for much of the spatial variation (Model 3): about 88% of the spatial variation (at the municipal level) is explained by cantonal differences and the considered covariates (bottom panel Fig 2).

**Fig 2.**
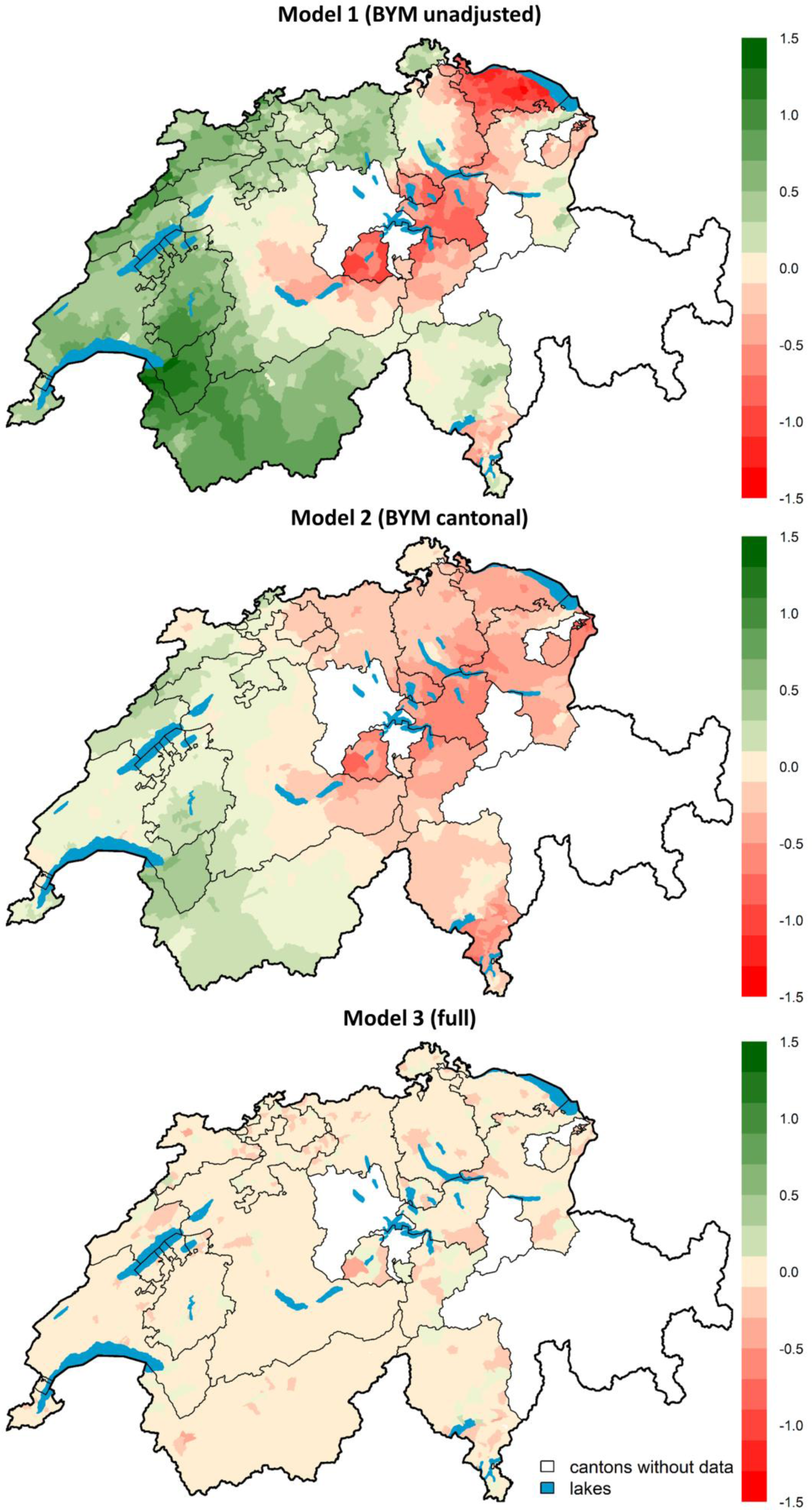
Spatial variation of HPV vaccination uptake in Switzerland at the municipal level. Top panel: spatial variation accounting only for correlation between neighbouring municipalities (model 1, Besag York Mollié model, BYM unadjusted); middle panel: remaining spatial variation after adjusting for cantonal differences (model 2, BYM cantonal); bottom panel: remaining spatial variation after adjusting for cantonal differences and covariates (model 3, full). Shown are the differences from the mean on the log odds scale. Municipalities with no information about HPV vaccination uptake borrow information from the first order neighbouring municipalities. White areas represent cantons for which we did not get authorization to analyse the data.

### Model selection

Of the fitted models 1-7, the model with the smallest DIC was the full model (model 3, DIC = 11,419) (Table 2). Results from the univariable models (model 8), the fully adjusted logit (model 7) and the full model (model 3) are shown in Fig 3 and in more detail in the online supplementary Table S3. The association of individual covariates (model 8) becomes weaker after adjusting for all covariates (model 7) and additionally including the random effects (model 3). We observed persistent strong associations for nationality, political opinion, availability of school-based HPV vaccination and survey period.

**Table 2.** Comparison of Bayesian hierarchical logistic regression models that explain the spatial heterogeneity of HPV vaccination uptake in Switzerland.

| Model type | Spatial component | Cantonal random effect | Covariates | DIC |
| --- | --- | --- | --- | --- |
| Model 1 (BYM unadjusted) | X |  |  | 11,513 |
| Model 2 (BYM cantonal) | X | X |  | 11,489 |
| Model 3 (full) | X | X | X | 11,417 |
| Model 4 | X |  | X | 11,432 |
| Model 5 |  | X |  | 11,541 |
| Model 6 |  | X | X | 11,450 |
| Model 7 (logit model) |  |  | X | 11,557 |
| Model 8 (univariable)* |  |  | X | - |
| Abbreviations: BYM Besag York Mollié prior, DIC Deviance Information Criterion<br>*Model 8 (univariable) was adjusted for one variable at time |  |  |  |  |

**Fig 3.**
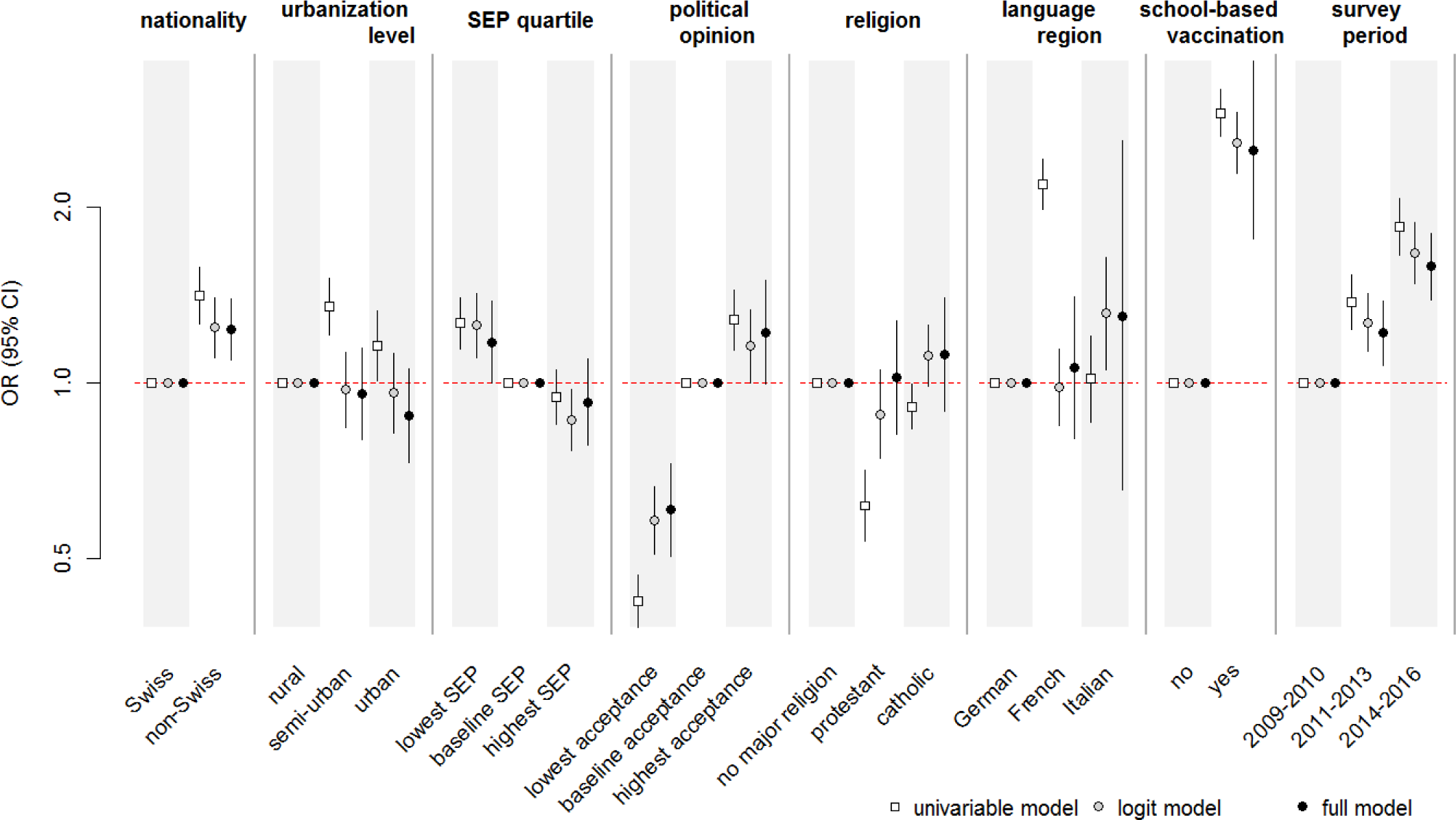
Odds ratios (OR) and 95% credible intervals for being vaccinated for HPV. The full model (model 3) is adjusted for all covariates and includes random effect terms to account for cantonal and municipal differences in uptake and spatial autocorrelation at the municipality-level. The adjusted-model (model 7) includes all covariates without any random effect terms. The univariable model (model 8), includes each covariate individually at a time without random effect terms.

### Associations between vaccination uptake and covariates

Availability of school-based vaccination delivery in a canton was strongly associated with higher vaccination uptake (Fig 3 and online supplementary Table S3, OR from full model: 2.51, 95% CI: 1.77-3.56). Living in a municipality in the lowest quartile of acceptance of the referendum on the revision of the epidemic law was associated with reduced uptake (OR: 0.61, 95% CI: 0.50-0.73) and living in a municipality in the highest quartile was associated with increased uptake (OR: 1.22, 95% CI: 0.99-1.50). These covariates were strongly correlated; only 40% of girls living in municipalities with low acceptance of the vote lived in a canton with school-based vaccination, compared with 85% of girls from municipalities with high acceptance (online supplementary Fig S8).

Vaccination uptake was higher in municipalities in the lowest Swiss-SEP quartile (OR: 1.18, 95% CI: 1.00-1.38) and among non-Swiss residents (OR: 1.23, 95% CI: 1.09-1.39). Vaccination uptake increased over the three survey periods (OR: 1.22, 95% CI: 1.07-1.38 and OR: 1.58, 95% CI: 1.38-1.81 for 2011-2013 and 2014-2016, respectively). In the full model, there was no evidence that uptake differed between high (highest quartile) and medium (2^nd^ and 3^rd^ quartile) levels of SEP (OR highest quartile: 0.93, 95% CI: 0.78-1.10).

In univariable models, living in a French-speaking municipality was associated with higher vaccination uptake while living in a rural or protestant municipality was associated with lower uptake (Fig 3 and online supplementary Table S3). However, there was little evidence that these factors were associated with HPV vaccine uptake after adjusting for other covariates and accounting for cantonal and municipal level differences (full model). Language region was highly correlated with both school-based vaccination (school-based vaccination was available in all French-speaking cantons) and vote results (<1% of girls from French-speaking regions lived in municipalities with lowest acceptance of the referendum, compared with 38% for German-speaking regions).

Our results did not change materially in sensitivity analyses. Imputing vaccine uptake in non-respondents using extreme assumptions (all vaccinated or all non-vaccinated) did not substantially change the estimated OR for most covariates (online supplementary material, Section 3 and Table S4-S5). The estimated ORs did change for nationality and survey period; non-Swiss individuals were over-represented among non-respondents, and the last survey period had a higher proportion of non-respondents compared to the other two survey periods. Accounting for different sampling methods in the full model resulted in similar estimates of OR for the covariates (online supplementary Table S5).

## Discussion

This spatial analysis of cross-sectional survey data found that cantonal differences and included covariates explained up to almost 90% of municipality-level variation in HPV vaccination uptake among girls in Switzerland. Availability of school-based vaccination delivery was strongly associated with increased HPV vaccination uptake and low municipal-level acceptance of a popular vote about the revision of the epidemic law, which included strengthening of vaccination promotion, was strongly associated with decreased uptake. Uptake of HPV vaccination increased over the three survey periods from 2009 to 2016.

The main strength of our study was the availability of survey data based on written vaccination records from cantons that cover more than 90% of the Swiss population during the evolution of HPV vaccination programs from 2009 until 2016. We were able to include a wide range of covariates at individual, municipal and cantonal level. Furthermore, our analysis accounted for spatial autocorrelation, which could result in inaccurate associations if ignored [28,46]. Our study has some limitations. First, the survey sampling methods and response rates differed between cantons. Our sensitivity analysis showed that the effect of covariates remained similar after accounting for differences in sampling methods. Second, our findings might have been affected by selection bias. If parents of vaccinated girls were more likely to respond to the survey, HPV vaccine uptake levels might be overestimated. However, even under extreme assumptions about vaccine uptake in non-respondents, the association of the covariates with vaccination uptake remained similar (except for nationality and survey period which had large differences in response rates). Third, the municipal-level covariates were based on data collected in the national census from 2000, before HPV vaccination programs began. We do not expect this to have affected our study because the composition of communities in Switzerland according to factors such as religion and socio-economic position show little variation over time [47]. Lastly, the most recent information about the organisation of cantonal HPV vaccination programmes, on which we based our analysis, was published in 2009. Some aspects of the programmes may have changed in some cantons since then.

To our knowledge, our study is the first analysis of spatial variation in HPV vaccination uptake in a country using outcome data at the individual level and adjusting for spatial autocorrelation. A systematic review of 25 studies of factors associated with HPV vaccine uptake, published up to 2011, found that most were cross-sectional studies from a single or limited number of states in the US [14]. The studies typically included factors at the individual level and controlled for no, or a limited number of potential confounders. The review highlighted that the highest levels of vaccination uptake came from studies with school-based programs, which corroborates our finding that availability of school-based vaccination delivery is associated with higher uptake. We found three studies that used spatial autocorrelation models, published since 2011, all from limited geographical areas in the US. The first targeted uninsured and publicly insured children in North Carolina [17], the second used data from an internet survey from the Twin Cities Metropolitan Area of Minnesota [22] and the third was based on seven Minnesota counties [16]. In these studies, substantial spatial variation remained after adjusting for their covariates. Our study covered the majority of a whole country and the full model explained almost all small-scale spatial variation. Two studies that examined geographical variation in HPV vaccination uptake according to voting patterns, in the Netherlands [11] and the US [15], found lower HPV vaccine uptake in areas that elected religious or conservative parties. An advantage of our study is that we used voting data from a referendum that was more closely linked to people’s attitudes towards vaccination than voting for a political party in general.

We found that girls living in municipalities with the lowest percentage acceptance of a vote to revise an epidemic law were less likely to be vaccinated. Some [18,20,21,27] but not all [11,16,19,20,23–26] studies have found an inverse association between SEP (or poverty based on income) and HPV vaccine uptake. This discrepancy between studies might be due to differences in national healthcare and health insurance systems. We found that girls living in municipalities in the lowest quartile of Swiss-SEP were more likely to receive HPV vaccination than in municipalities in the middle quartiles in the univariable, but not the multivariable analysis. Our finding that non-Swiss girls were more likely to be vaccinated than Swiss girls could mean that people with family origins outside Switzerland accept vaccination more readily than Swiss people but it might also reflect household-level socio-economic disparities that were not captured by the Swiss-SEP. Since non-Swiss girls had a higher non-response rate, interpretation should be treated with caution. Ethnicity was not recorded in the SNVCS and is not routinely recorded in Switzerland. Other studies that have considered ethnicity report lower levels of HPV vaccine uptake in girls from non-white ethnic groups [14].

The findings of our study support the hypothesis that there is interplay between people’s attitudes about vaccination, the availability of vaccination services and the probability of an individual girl receiving HPV vaccination. The best-fitting model included a random effect at the cantonal level and, together with selected covariates, explained almost all small-scale spatial variation in HPV vaccine uptake. Cantons have considerable autonomy in providing health services and represent a contextual factor for vaccine accessibility. Vaccine scepticism in a community, in turn, could impact the political outcome of decision makers and hence affect vaccination policies. Our findings do not necessarily represent causal associations because of the ecological nature of the associations and the cross-sectional nature of our study design. The strong association between HPV vaccination uptake and patterns of voting about vaccination laws at the municipality level are, however, consistent with the suggestion that scepticism or opposition to vaccination could influence decisions of parents and their daughters to get vaccinated. A nationally representative Swiss survey found that fear of side effects and general opposition to vaccination were two of the main reasons that participants gave for not being vaccinated against HPV [48]. The vote results were also strongly correlated with the availability of school-based vaccination. This relation might indicate how individuals shape their community, thereby influencing health services and affecting health outcomes. Thus, the difficulty to achieve higher levels of HPV vaccination uptake in areas with high levels of vaccine scepticism result not only from vaccine refusal, but also from a lack of easy access to vaccination such as through school-based delivery.

We conducted a multi-level spatial analysis to identify the factors that are associated with the spatial variation in HPV vaccination uptake in Switzerland. Our study highlights the importance of taking into account spatial autocorrelation and covariates at different spatial levels. Our results support the importance of an interplay between regional contextual factors and vaccine scepticism in determining HPV vaccination uptake. Our study suggests that higher levels of HPV vaccination could be achieved by efforts to mitigate vaccine scepticism, which might then permit broader use of school-based delivery of HPV vaccination.

## Acknowledgements

We would like to thanks the cantons for providing access to their HPV vaccination survey data. We would also like to thank Dr. Jan von Overbeck and Vanessa Arn for their help in obtaining agreements from the cantons.

## Conflict of interest

The authors declare that they have no conflict of interest.

## Research ethics statement

Permission for the use of anonymised data from the SNVCS was obtained for all participating cantons. Ethical approval was not required because we used anonymised health-related data (according to the Swiss Human Research Act (HRA, Art.2.2 al.c.).

## Author contributions

Conceptualisation, M.R., G.K.¸ C.L.A., B.S., N.L. and M.B.; Methodology, G.K., M.R., B.S., C.L.A. and N.L.; Formal analysis, G.K; Validation, B.S, N.L, C.L.A, C.H, P.L, A.S, M.B and M.M; Writing-original draft, M.R. and G.K; Writing – review and editing, M.R., G.K., C.L.A., B.S., N.L., P.L., A.S., M.M., M.B. and C.H.; Resources, P.L., C.H., A.S. and M.M.; Supervision, C.L.A, B.S and N.L.

## Funding

This study was supported by the Swiss Cancer League and the Swiss Cancer Research foundation (Grant No. 3049-08-2012 and 3515-08-2014). B.D. Spycher was supported by a Swiss National Science Foundation fellowship (PZ00P3_147987).

## References

1. World Health Organization W. Human papillomavirus (HPV) and cervical cancer [Internet]. Fact sheet; 2016. Available: http://www.who.int/mediacentre/factsheets/fs380/en/

2. Bruni L, Diaz M, Barrionuevo-Rosas L, Herrero R, Bray F, Bosch FX, et al. Global estimates of human papillomavirus vaccination coverage by region and income level: a pooled analysis. Lancet Glob Health. 2016;4: e453–e463. doi:10.1016/S2214-109X(16)30099-7

3. Brisson M, van de Velde N, Franco EL, Drolet M, Boily M-C. Incremental Impact of Adding Boys to Current Human Papillomavirus Vaccination Programs: Role of Herd Immunity. J Infect Dis. 2011;204: 372–376. doi:10.1093/infdis/jir285

4. Public Health England. Human Papillomavirus (HPV) vaccination coverage in adolescent females in England: 2015/2016 [Internet]. Dec 2016 [cited 29 Jun 2017]. Available: https://www.gov.uk/government/uploads/system/uploads/attachment_data/file/578729/HPV_vaccination-_2015-16.pdf

5. National HPV Register. HPV Vaccination Coverage 2015 - National Australia HPV Vaccination Program Register for females turning 15 years of age in 2015 [Internet]. Feb 2017 [cited 29 Jun 2017]. Available: http://www.hpvregister.org.au/research/coverage-data/HPV-Vaccination-Coverage-2015

6. Potts A, Sinka K, Love J, Gordon R, McLean S, Malcolm W, et al. High uptake of HPV immunisation in Scotland – perspectives on maximising uptake. 18. 2013;39. Available: http://www.eurosurveillance.org/ViewArticle.aspx?ArticleId=20593

7. Giambi C, Donati S, Declich S, Salmaso S, degli Atti MLC, Alibrandi MP, et al. Estimated acceptance of HPV vaccination among Italian women aged 18–26 years. Vaccine. 2011;29: 8373–8380. doi:10.1016/j.vaccine.2011.08.079

8. Héquet D, Rouzier R. Determinants of geographic inequalities in HPV vaccination in the most populated region of France. PLOS ONE. 2017;12: e0172906. doi:10.1371/journal.pone.0172906

9. Poethko-Müller C, Buttmann-Schweiger N, KiGGS Study Group. [HPV vaccination coverage in German girls: results of the KiGGS study: first follow-up (KiGGS Wave 1)]. Bundesgesundheitsblatt Gesundheitsforschung Gesundheitsschutz. 2014;57: 869–877. doi:10.1007/s00103-014-1987-3

10. Durham DP, Ndeffo-Mbah ML, Skrip LA, Jones FK, Bauch CT, Galvani AP. National- and state-level impact and cost-effectiveness of nonavalent HPV vaccination in the United States. Proc Natl Acad Sci. 2016;113: 5107–5112. doi:10.1073/pnas.1515528113

11. Rondy M, van Lier A, van de Kassteele J, Rust L, de Melker H. Determinants for HPV vaccine uptake in the Netherlands: A multilevel study. Vaccine. 2010;28: 2070–2075. doi:10.1016/j.vaccine.2009.12.042

12. Holman DM, Benard V, Roland KB, Watson M, Liddon N, Stokley S. Barriers to human papillomavirus vaccination among US adolescents: a systematic review of the literature. JAMA Pediatr. 2014;168: 76–82. doi:10.1001/jamapediatrics.2013.2752

13. Bühlmann M. Politische Partizipation im kommunalen Kontext: der Einfluss lokaler Kontexteigenschaften auf individuelles politisches Partizipationsverhalten. 1. Aufl. Bern: Haupt; 2006.

14. Kessels SJM, Marshall HS, Watson M, Braunack-Mayer AJ, Reuzel R, Tooher RL. Factors associated with HPV vaccine uptake in teenage girls: a systematic review. Vaccine. 2012;30: 3546–3556. doi:10.1016/j.vaccine.2012.03.063

15. Bernstein S, North A, Schwartz J, Niccolai LM. State-Level Voting Patterns and Adolescent Vaccination Coverage in the United States, 2014. Am J Public Health. 2016;106: 1879–1881. doi:10.2105/AJPH.2016.303381

16. Finney Rutten LJ, Wilson PM, Jacobson DJ, Agunwamba AA, Radecki Breitkopf C, Jacobson RM, et al. A Population-Based Study of Sociodemographic and Geographic Variation in HPV Vaccination. Cancer Epidemiol Biomark Prev Publ Am Assoc Cancer Res Cosponsored Am Soc Prev Oncol. 2017;26: 533–540. doi:10.1158/1055-9965.EPI-16-0877

17. Trogdon JG, Ahn T. Geospatial Patterns in Human Papillomavirus Vaccination Uptake: Evidence from Uninsured and Publicly Insured Children in North Carolina. Cancer Epidemiol Biomarkers Prev. 2015;24: 595–602. doi:10.1158/1055-9965.EPI-14-1231

18. Henry KA, Stroup AM, Warner EL, Kepka D. Geographic Factors and Human Papillomavirus (HPV) Vaccination Initiation among Adolescent Girls in the United States. Cancer Epidemiol Biomark Prev Publ Am Assoc Cancer Res Cosponsored Am Soc Prev Oncol. 2016;25: 309–317. doi:10.1158/1055-9965.EPI-15-0658

19. Guthmann J-P, Pelat C, Célant N, Parent du Chatelet I, Duport N, Rochereau T, et al. Socioeconomic inequalities to accessing vaccination against human papillomavirus in France: Results of the Health, Health Care and Insurance Survey, 2012. Rev Epidemiol Sante Publique. 2017;65: 109–117. doi:10.1016/j.respe.2017.01.100

20. Pruitt SL, Schootman M. Geographic disparity, area poverty, and human papillomavirus vaccination. Am J Prev Med. 2010;38: 525–533. doi:10.1016/j.amepre.2010.01.018

21. Tsui J, Gee GC, Rodriguez H, Kominski GF, Glenn BA, Singhal R, et al. Exploring the role of neighborhood socio-demographic factors on HPV vaccine initiation among low-income, ethnic minority girls. J Immigr Minor Health Cent Minor Public Health. 2013;15: 732–740. doi:10.1007/s10903-012-9736-x

22. Nelson EJ, Hughes J, Oakes JM, Pankow JS, Kulasingam SL. Geospatial patterns of human papillomavirus vaccine uptake in Minnesota. Bmj Open. 2015;5: e008617. doi:10.1136/bmjopen-2015-008617

23. Lefevere E, Hens N, De Smet F, Van Damme P. Dynamics of HPV vaccination initiation in Flanders (Belgium) 2007-2009: a Cox regression model. BMC Public Health. 2011;11: 470. doi:10.1186/1471-2458-11-470

24. Smith LM, Brassard P, Kwong JC, Deeks SL, Ellis AK, Lévesque LE. Factors associated with initiation and completion of the quadrivalent human papillomavirus vaccine series in an ontario cohort of grade 8 girls. BMC Public Health. 2011;11: 645. doi:10.1186/1471-2458-11-645

25. Schülein S, Taylor KJ, König J, Claus M, Blettner M, Klug SJ. Factors influencing uptake of HPV vaccination among girls in Germany. BMC Public Health. 2016;16: 995. doi:10.1186/s12889-016-3663-z

26. Centers for Disease Control and Prevention (CDC). National, state, and local area vaccination coverage among adolescents aged 13-17 years --- United States, 2009. MMWR Morb Mortal Wkly Rep. 2010;59: 1018–1023.

27. Gerend MA, Weibley E, Bland H. Parental response to human papillomavirus vaccine availability: uptake and intentions. J Adolesc Health Off Publ Soc Adolesc Med. 2009;45: 528–531. doi:10.1016/j.jadohealth.2009.02.006

28. Langford IH, Leyland AH, Rasbash J, Goldstein H. Multilevel modelling of the geographical distributions of diseases. J R Stat Soc Ser C Appl Stat. 1999;48: 253–268.

29. Bundesamt für Gesundheit. Vaccination contre le cancer du col de l’utérus-Début des programmes cantonaux de vaccination. Bulletin 38; 2008.

30. Spaar A, Masserey V. HPV-Impfung in der Schweiz. Impfraten und aktuelle Daten zu Wirksamkeit und Sicherheit. Pädiatrie. 2015;6:8–11.

31. Bundesamt für Gesundheit. Die HPV-Impfprogramme in der Schweiz: eine Synthese von 2007 bis 2010. Bulletin. 43:949–53. Available: https://www.bag.admin.ch/dam/bag/de/dokumente/mt/infektionskrankheiten/hpv/hpv-artikel-3.pdf.download.pdf/hpv-artikel-3-de.pdf

32. EBPI-Universität Zürich, Bundesamt für Gesundheit. Durchimpfung von 2-, 8- und 16- Jährigen Kindern in der Schweiz, 1999-2016 [Internet]. Available: http://www.bag.admin.ch/themen/medizin/00682/00685/02133/index.html?lang=de

33. Bundesamt für Gesundheit. Durchimpfung von 2-,8- und 16-Jährigen in der Schweiz, 2011 bis 2013. Bulletin. 2015;28.

34. Bundesamt für Statistik. Statistik der Bevölkerung und der Haushalte (STATPOP), Sektion Demografie und Migration. In: Ständige und nichtständige Wohnbevölkerung nach institutionellen Gliederungen, Staatsangehörigkeit (Kategorie), Geschlecht und Alter [Internet]. 2013 [cited 12 Sep 2017]. Available: https://www.bfs.admin.ch/bfs/de/home/statistiken/bevoelkerung/migration-integration/auslaendische-bevoelkerung.assetdetail.3262116.html

35. PLANeS, Schweizerische Stiftung für sexuelle un reproduktive Gesundheit. Evaluation kantonaler HPV-Impfprogramme, Definitiver Bericht für das BAG. Bern; 2009 p. 21. Report No.: 09.002385.

36. Bundesamt für Statisik. Die Raumgliederungen der Schweiz 2016 - MS-Excel Version [Internet]. Bundesamt für Statistik, Schweizerische Eidgenossenschaft; 2017 Mar. Report No.: be-b-00.04-rgs-16. Available: https://www.bfs.admin.ch/bfs/de/home/grundlagen/raumgliederungen.assetdetail.2118475.html

37. Bundesamt für Statistik. Swiss Population Census 2000 [Internet]. Bern, Schweiz; 2000. Available: https://www.bfs.admin.ch/bfs/en/home/statistics/population/surveys/census.html

38. Panczak R, Galobardes B, Voorpostel M, Spoerri A, Zwahlen M, Egger M, et al. A Swiss neighbourhood index of socioeconomic position: development and association with mortality. J Epidemiol Community Health. 2012;66: 1129–1136. doi:10.1136/jech-2011-200699

39. Panczak R, Zwahlen M, Woitek U, Rühli FJ, Staub K. Socioeconomic, Temporal and Regional Variation in Body Mass Index among 188,537 Swiss Male Conscripts Born between 1986 and 1992. PLOS ONE. 2014;9: e96721. doi:10.1371/journal.pone.0096721

40. Federal Chancellery, Swiss Confederation. Loi fédérale sur la lutte contre les maladies transmissibles de l’homme (Loi sur les épidémies, LEp) [Internet]. [cited 19 May 2017]. Available: https://www.admin.ch/ch/f/pore/rf/cr/2007/20071012.html

41. Bundesamt für Statistik. Statistik der Wahlen und Abstimmungen, Abstimmungen 2013: Epidemiengesetz [Internet]. Neuchâtel, Suisse; 2013 Sep. Available: https://www.bfs.admin.ch/bfs/de/home/statistiken/politik/erhebungen/swa.html

42. Maheswaran R, Haining RP, Pearson T, Law J, Brindley P, Best NG. Outdoor NOx and stroke mortality: adjusting for small area level smoking prevalence using a Bayesian approach. Stat Methods Med Res. 2006;15: 499–516. doi:10.1177/0962280206071644

43. Panczak R, Held L, Moser A, Jones PA, Rühli FJ, Staub K. Finding big shots: small-area mapping and spatial modelling of obesity among Swiss male conscripts. BMC Obes. 2016;3. doi:10.1186/s40608-016-0092-6

44. Spiegelhalter DJ, Best NG, Carlin BP, Van Der Linde A. Bayesian measures of model complexity and fit. J R Stat Soc Ser B Stat Methodol. 2002;64: 583–639. doi:10.1111/1467-9868.00353

45. Rue H, Martino S, Chopin N. Approximate Bayesian inference for latent Gaussian models by using integrated nested Laplace approximations. J R Stat Soc Ser B Stat Methodol. 2009;71: 319–392. doi:10.1111/j.1467-9868.2008.00700.x

46. Werneck GL, Maguire JH. Spatial modeling using mixed models: an ecologic study of visceral leishmaniasis in Teresina, Piauí State, Brazil. Cad Saúde Pública. 2002;18: 633–637. doi:10.1590/S0102-311X2002000300007

47. Bundesamt für Statistik. Religions. In: Religionen [Internet]. [cited 5 Sep 2017]. Available: https://www.bfs.admin.ch/bfs/fr/home/statistiques/population/langues-religions/religions.html

48. Bundesamt für Gesundheit, Abteilung übertragbare Krankheiten. Die HPV-Impfung in der Schweiz: Resulate einer nationalen Befragung im Jahr 2014. Bulletin. 2015;23/15: 445–452.

